# Household-Level Risk Factors for *Aedes aegypti* Pupal Density in Guayaquil, Ecuador

**DOI:** 10.1101/2020.11.23.391938

**Authors:** Thien-An Ha, Tomás M. León, Karina Lalangui, Patricio Ponce, John M. Marshall, Varsovia Cevallos

**Affiliations:** School of Public Health, University of California, Berkeley, USA; Centro de Investigación en Vectores Artrópodos, Instituto Nacional de Investigación en Salud Pública “Dr. Leopoldo Izquieta Pérez”, Quito, Ecuador

**Keywords:** aedes aegypti, mosquito, household risk factors, arbovirus, collection services, precipitation, predictive modeling

## Abstract

**Background:** Vector-borne diseases are a major cause of disease burden in Guayaquil, Ecuador, especially arboviruses spread by *Aedes aegypti* mosquitoes. Understanding which household characteristics and risk factors lead to higher *Ae. aegypti* densities and consequent disease risk can help inform and optimize vector control programs.

**Methods:** Cross-sectional entomological surveys were conducted in Guayaquil between 2013 and 2016, covering household demographics, municipal services, potential breeding containers, presence of *Ae. aegypti* larvae and pupae, and history of using mosquito control methods. A zero-truncated negative binomial regression model was fitted to data for estimating the household pupal index. An additional model assessed the factors of the most productive breeding sites across all of the households.

**Results:** Of surveyed households, 610 satisfied inclusion criteria. The final household-level model found that collection of large solid items (e.g., furniture and tires) and rainfall the week of and 2 weeks before collection were negatively correlated with average pupae per container, while bed canopy use, unemployment, container water volume, and the interaction between large solid collection and rainfall 2 weeks before the sampling event were positively correlated. Selection of these variables across other top candidate models with ΔAICc < 1 was robust, with the strongest effects from large solid collection and bed canopy use. The final container-level model explaining the characteristics of breeding sites found that contaminated water is positively correlated with *Ae. aegypti* pupae counts while breeding sites composed of car parts, furniture, sewerage parts, vases, ceramic material, glass material, metal material, and plastic material were all negatively correlated.

**Conclusion:** Having access to municipal services like bulky item pickup was effective at reducing mosquito proliferation in households. Association of bed canopy use with higher mosquito densities is unexpected, and may be a consequence of large local mosquito populations or due to limited use or effectiveness of other vector control methods. The impact of rainfall on mosquito density is multifaceted, as it may both create new habitat and “wash out” existing habitat. Providing services and social/technical interventions focused on monitoring and eliminating productive breeding sites is important for reducing aquatic-stage mosquito densities in households at risk for *Ae. aegypti-transmitted* diseases.

## Background

Vector-borne febrile illnesses such as dengue, chikungunya, and Zika virus are of pressing public health concern in Latin America and the Caribbean [1]. The mosquito *Aedes aegypti* is the region’s primary vector of these arboviruses, which co-circulate in populations in the tropics and subtropics [1,2]. The burden of these diseases weighs heavily on susceptible populations in low and middle-income countries such as Ecuador [1].

Between 2010 and 2014, over 70,000 cases of dengue were reported in Ecuador, with the highest incidence clustered in urbanized coastal areas like the city of Guayaquil [1,3]. Dengue infection can be asymptomatic or present as a moderate febrile illness, with some symptoms advancing to hemorrhage, shock, and death [4]. Without an effective dengue vaccine, community and household-level vector control of *Ae. aegypti* remains the primary means of preventing and controlling dengue outbreaks [2]. In Ecuador, each household currently employs, on average, five different mosquito control methods, including sprays, aerosols, repellents, mosquito coils, screens, and bed nets [1].

*Ae. aegypti* population management is an ongoing public health challenge for countries with limited resources that must efficiently plan and utilize targeted control. *Ae. aegypti* is a mosquito species that primarily amplifies epidemics among urban populations [5]. The species is an effective vector for dengue because it is highly adapted to urban environments, where it lays eggs in artificial containers of water near human dwellings and preferentially feeds on humans [6]. Adult *Ae. aegypti* lay eggs in such habitats, and larvae develop in both natural water-retaining structures and in domestic water containers [7].Examples of outdoor breeding sites for *Ae. aegypti* include large tires, flower vases, and plastic gallon containers [8]. Understanding the local characteristics of *Ae. aegypti* habitats can be used to inform vector control efforts [9]. Previous studies done in Machala, Ecuador, found that local socio-ecological conditions such as proximity to abandoned properties, interruptions in the piped water supply, and a highly shaded patio were risk factors for *Ae. aegypti* proliferation and the presence of dengue [2]. Further investigation into household factors, in conjunction with the evaluation of vector control efforts, is necessary to reduce and prevent dengue incidence by reducing *Ae. aegypti* habitat and population.

Our study describes potential household-level risk factors for *Ae. aegypti* pupal proliferation in the city of Guayaquil, Ecuador’s largest and most populous city, its most important commercial port, and the historical epicenter of yellow fever and dengue in the country[1]. Guayaquil (2016 Instituto Nacional de Investigación en Salud Pública projected population: 2,482,789), is located on the west bank of the Guayas River (Fig. 1), which flows into the Pacific Ocean. The urban core of Guayaquil is surrounded by low-income neighborhoods with limited basic services and high rates of migration. Guayaquil has had the greatest number of dengue cases in Ecuador since 1988, with all 4 serotypes circulating since then. Seasonally, the highest incidence occurs in the rainy season because of favorable environmental conditions for transmission. The first four months of the year have abundant rain in this coastal region, with over 17 average days of rain totalling more than 200 mm each month; in the province of Guayas, the days are hot and humid, with average high temperatures between 29°C and 32°C and high humidity (NOAA ASOS Data).

**Fig 1.**
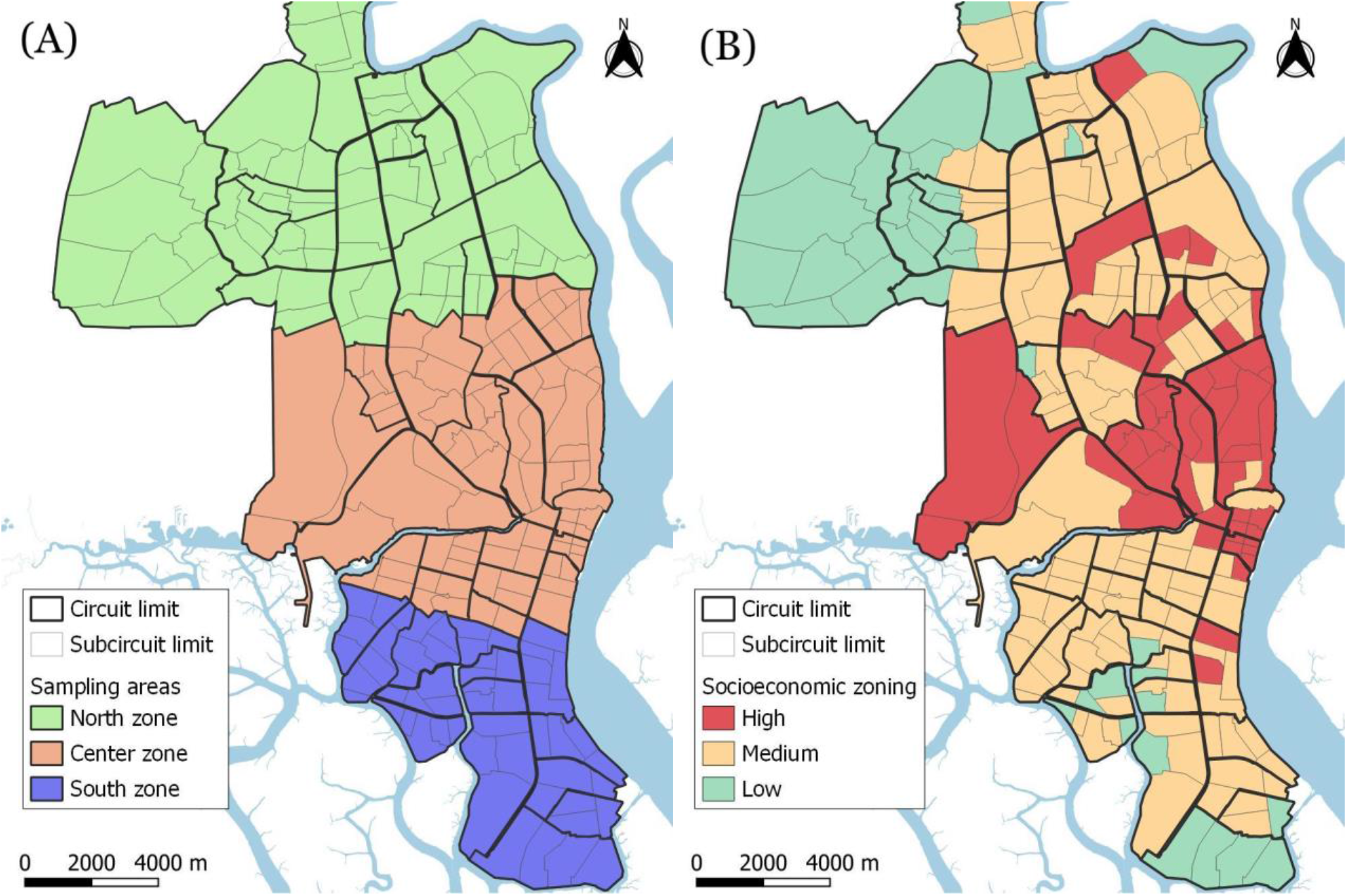
Subcircuits and zones of Guayaquil showing sampling zones (A) and socioeconomic status (B) by subcircuit.

## Methods

### Household Data Collection

Household cross-sectional surveys were conducted every month in Guayaquil from January 2013 to August 2016 to investigate factors correlated with mosquito pupal counts per container. Each month, one random subcircuit (an administrative unit covering ~10,000 inhabitants) was randomly selected in the North, Center, and South of Guayaquil, resulting in three sampling events (Fig. 1A). Guayaquil subcircuits were also classified into low, medium, and high socioeconomic status derived from three indicators: illiteracy, overcrowding, and unemployment (Fig. 1B). Each of these variables were averaged across the subcircuit, where illiteracy and unemployment were represented as percentages and overcrowding was the average number of people per household.

Households were defined as in-use residential units. Household addresses were obtained prior to the survey. Containers were defined as in-use breeding sites near or inside the household. Each container was examined for immature mosquito stages in the water, material, and type. Each house was visited only once in order to maximize the geographical area covered. Each visit assessed mosquito presence and conducted sampling in artificial containers of water. During each house visit, questions were asked to any household member over the age of 18 after verifying their residency. These questions focused on *Ae. aegypti* risk factors, including Ministry of Health vector control efforts and mosquito control practices which were used as candidate predictors in our model (Table 1, Table S1).

**Table 1.**
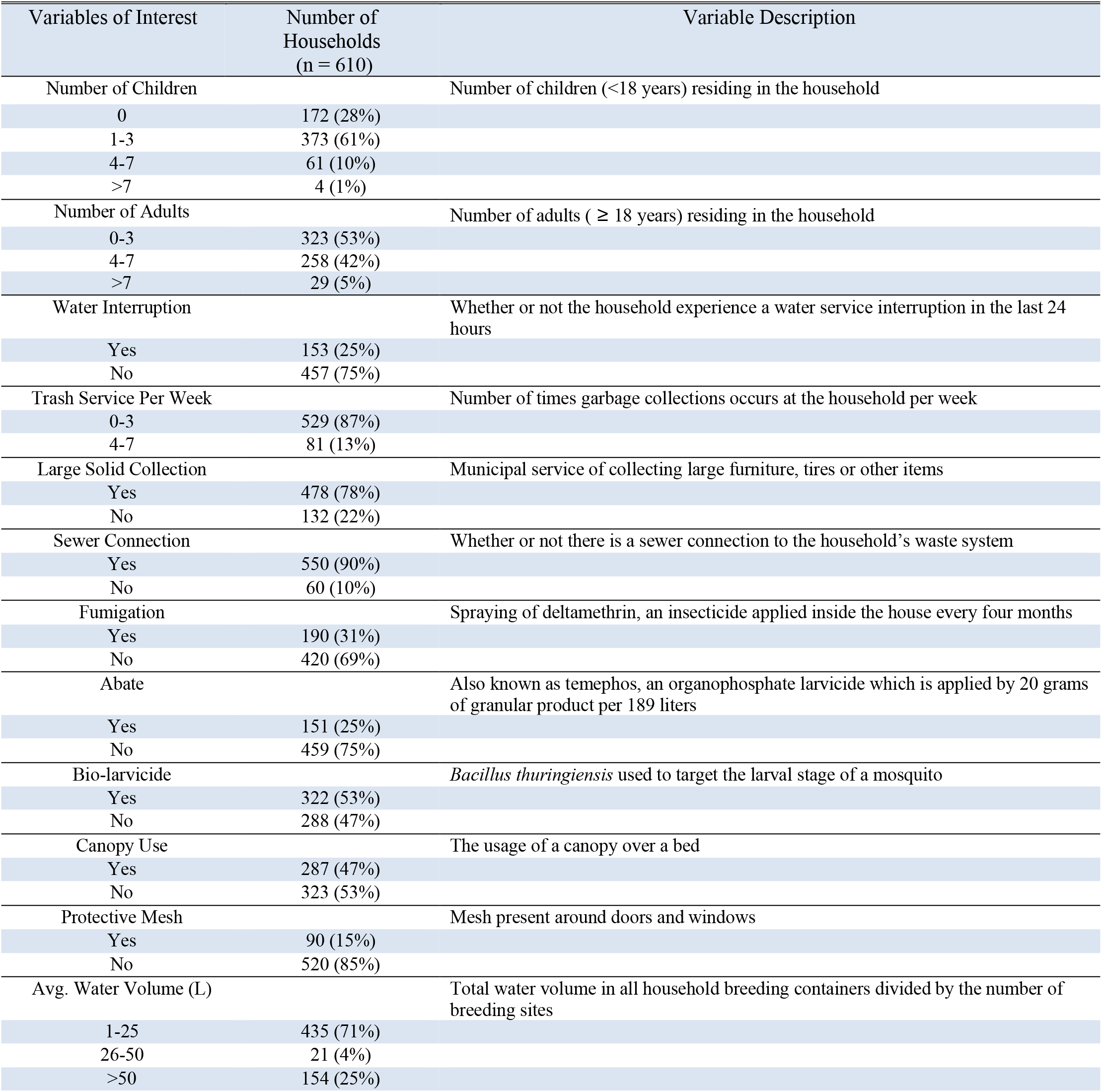
Summary statistics for habitat and vector control effort variables included in the analysis.

| Variables of Interest | Number of Households (n = 610) | Variable Description |
| --- | --- | --- |
| Number of Children |  | Number of children (<18 years) residing in the household |
| 0 | 172 (28%) |  |
| 1-3 | 373 (61%) |  |
| 4-7 | 61 (10%) |  |
| >7 | 4 (1%) |  |
| Number of Adults |  | Number of adults ( ≥ 18 years) residing in the household |
| 0-3 | 323 (53%) |  |
| 4-7 | 258 (42%) |  |
| >7 | 29 (5%) |  |
| Water Interruption |  | Whether or not the household experience a water service interruption in the last 24 hours |
| Yes | 153 (25%) |  |
| No | 457 (75%) |  |
| Trash Service Per Week |  | Number of times garbage collections occurs at the household per week |
| 0-3 | 529 (87%) |  |
| 4-7 | 81 (13%) |  |
| Large Solid Collection |  | Municipal service of collecting large furniture, tires or other items |
| Yes | 478 (78%) |  |
| No | 132 (22%) |  |
| Sewer Connection |  | Whether or not there is a sewer connection to the household's waste system |
| Yes | 550 (90%) |  |
| No | 60 (10%) |  |
| Fumigation |  | Spraying of deltamethrin, an insecticide applied inside the house every four months |
| Yes | 190 (31%) |  |
| No | 420 (69%) |  |
| Abate |  | Also known as temephos, an organophosphate larvicide which is applied by 20 grams of granular product per 189 liters |
| Yes | 151 (25%) |  |
| No | 459 (75%) |  |
| Bio-larvicide |  | *Bacillus thuringiensis* used to target the larval stage of a mosquito |
| Yes | 322 (53%) |  |
| No | 288 (47%) |  |
| Canopy Use |  | The usage of a canopy over a bed |
| Yes | 287 (47%) |  |
| No | 323 (53%) |  |
| Protective Mesh |  | Mesh present around doors and windows |
| Yes | 90 (15%) |  |
| No | 520 (85%) |  |
| Avg. Water Volume (L) |  | Total water volume in all household breeding containers divided by the number of breeding sites |
| 1-25 | 435 (71%) |  |
| 26-50 | 21 (4%) |  |
| >50 | 154 (25%) |  |

Water interruptions were characterized by whether or not the household experienced a water service interruption in the last 24 hours. Collection of large solids are the municipal service of collecting large furniture or disposed tires. Adult mosquito fumigation, Abate (temephos), and biolarvicide are Ministry of Health vector control efforts that were implemented between 2013 and 2016. These vector control efforts were self-reported by the household member for previous weeks up to a month. Adult mosquito fumigation refers to the spraying of deltamethrin, an insecticide that is applied inside the house every four months [10]. The Ministry of Health decided upon which houses to fumigate based on historical locations of high numbers of dengue cases. Abate, the commercial name for temephos, is an organophosphate larvicide which is applied following World Health Organization (WHO) guidelines of 20g of granular product per 189 liters [10]. This product is applied to water containers 3 to 4 times a year [10], and is commonly used as a dengue vector control method [11]. The Ministry of Health distribution of abate was chosen based on previous larvae indices. Biolarvicide refers to the use of organisms such as bacteria to target the larval stage of a mosquito. In Guayaquil, the Ministry of Health applied and distributed the biolarvicide *Bacillus thuringiensis israelensis*. Protective mesh refers to whether the household has mesh around windows and doors.

Precipitation measurements at week 0, week 1 lag, and week 2 lag were also included. Week 0 indicates the amount of rainfall the week that the sampling event occurred. The lag variables, week 1 lag and week 2 lag, indicate precipitation one week previous to the sampling event and precipitation two weeks previous to the sampling event, respectively. Only variables that were most relevant to mosquito ecology based on literature review and entomologist consultation were included for the models.

### Entomological Data

Three field technicians conducted each sampling event, which took place across 250 households over a period of five days each time. In each house, technicians searched for immature mosquitoes in containers both inside and outside of the household. Each container carrying immatures was recorded for its container type and material type. These live immatures were transported to an insectary in Quito, where the lab at INSPI recorded the numbers and stage of development. Immatures were reared until adult stage for full species identification.

### Statistical Analysis

All analyses were performed in R version 3.5.3. We used zero-truncated negative binomial models to assess the appropriate household-level and container-level predictors of *Ae. aegypti* pupal population. Zero-truncated models were used because the data was only recorded for containers with present immature mosquitoes. Our outcome for the household-level statistical models was average pupal counts per container (APC), or pupal index, and was calculated as the total number of household pupae divided by the number of containers carrying these immature stage mosquitoes. Our outcome for the secondary analysis on container-level data was the sum of the pupae in each artificial breeding site. We determined that outliers for each outcome were those beyond Q3 (75th percentile) + 1.5 * IQR (Q3 - Q1) and omitted them [12].

We performed a chi-squared test for the Poisson model assumption that conditional variance equals conditional mean in our data set. We rejected the null hypothesis that the Poisson model best fits our data (p < 0.01) and fitted a negative binomial model with a dispersion parameter of 1.4103 and a standard error of 0.0881. We fitted a full model using all variables for the household model (Table S1) and the container model (Table S2) and used the ‘dredge’ function (R package MuMIn v1.43.17) to find all possible models through the best subset selection technique [13]. Best subset selection exhaustively searches all combinations of candidate variables and ranks models using specific selection criteria. In this study, we used the Akaike information criterion corrected for small sample sizes (AICc) to compare all the candidate models (Fig S1). AICc has a penalty term for small sample sizes: as sample size increases, AIC is approximated and therefore, AICc is preferred over AIC [14]. Effect sizes were considered significant if 95% confidence intervals for corresponding explanatory variables did not overlap zero. We evaluated model performances using 100 simulations of 10-fold cross-validation as our model evaluation method.

## Results

In the surveys, 830 households of the total 990 households surveyed were found to have *Ae. aegypti* mosquito pupae. Of the 830 households, 220 had missing or erroneous location data (e.g., coordinates indicated a household was not in Guayaquil) and were omitted, yielding 610 households for our analysis. The mean pupal index for the subset of included households was 11.08, with a standard deviation of 10.26, and a maximum pupal index of 42.

About 47% of the 610 households used bed canopies as a method for preventing mosquito biting (Table 1). Approximately 25% of households had water service interruptions. Only 22% of households did not have large solid collection services. About 15% of households had protective mesh around their windows and doors.

### Influence of household-related factors on Ae. aegypti pupal abundance

The model with the smallest (best) AICc value included the variables: canopy use, large solid service, unemployment, water volume, precipitation at week 0, precipitation at week 2 lag, and the interaction of large solid service and precipitation at week 2 lag (Table 2). The predictors with statistically significant associations were consistently selected into our top models ΔAICc < 1 (Table 2). The explanatory variable estimates for the model with the smallest AICc value, our top model, indicated that canopy use, unemployment, average container water volume, and the interaction between large solid service and precipitation (with 2-week lag) all had a statistically significant positive relationship with *Ae. aegypti* pupal abundance and large solid service had a significant negative relationship with *Ae. aegypti* pupal abundance (Table 3).

**Table 2.** Variables included in models with the smallest (best) AICc values. Only models that lie within a ΔAICc of 1 of the smallest AICc value are shown. The response variable was the total number of household *Ae. aegypti* pupae over the total number of household breeding sites. Error indicates the cross-validation prediction error off of the mean pupal index of 11.08. Precipitation0 indicates rainfall from the week of sampling and Precipitation2 indicates rainfall with 2-week lag (i.e., 2 weeks before sampling).

| Model | df | LogLik | $\Delta AICc$ | Error |
| --- | --- | --- | --- | --- |
| Canopy Use + Large Solid Service + Unemployment + Water Volume + Precipitation<sub>0</sub> + Precipitation<sub>2</sub> + Large Solid Service * Precipitation<sub>2</sub> | 9 | -2075.162 | 0 | 10.08 |
| Canopy Use + Water Interruption + Large Solid Service + Unemployment + Water Volume + Precipitation<sub>0</sub> + Precipitation<sub>2</sub> + Water Interruption * Precipitation<sub>0</sub> + Large Solid Service * Precipitation<sub>2</sub> | 11 | -2073.210 | 0.2374 | 10.12 |
| Canopy Use + Large Solid Service + Unemployment + Water Volume + Precipitation<sub>2</sub> + Large Solid Service * Precipitation<sub>2</sub> | 8 | -2076.444 | 0.5036 | 10.10 |
| Biolarvicide + Canopy Use + Large Solid Service + Unemployment + Water Volume + Precipitation<sub>0</sub> + Precipitation<sub>2</sub> + Large Solid Service * Precipitation<sub>2</sub> | 10 | -2074.407 | 0.5566 | 10.09 |
| Biolarvicide + Canopy Use + Water Interruption + Large Solid Service + Unemployment + Water Volume + Precipitation<sub>0</sub> + Precipitation<sub>2</sub> + Water Interruption * Precipitation<sub>0</sub> + Large Solid Service * Precipitation<sub>2</sub> | 12 | -2072.431 | 0.7603 | 10.13 |

**Table 3.** Explanatory variable estimates for the model with the smallest AICc value. The response variable is the average pupae per container in a household.

| Variable | Log Estimate | Estimate | 95% CI |
| --- | --- | --- | --- |
| Intercept | 1.790 | 6.00 | (3.68, 9.79) |
| Canopy Use | 0.240 | 1.271* | (1.06, 1.52) |
| Large Solid Service | -0.280 | 0.756* | (0.610, 0.932) |
| Unemployment | 0.0641 | 1.0662* | (1.02, 1.114) |
| Water Volume | 0.00166 | 1.0016* | (1.0003, 1.003) |
| Precipitation<sub>0</sub> | -0.00600 | 0.994 | (0.986, 1.0013) |
| Precipitation<sub>2</sub> | -0.0103 | 0.990 | (0.970, 1.0118) |
| Large Solid Service * Precipitation<sub>2</sub> | 0.0214 | 1.0216* | (1.00, 1.043) |

\*p < 0.05

The average prediction error over 100 simulations of 10-fold cross-validation was 10 pupae off of the true value where the mean pupal index was 11.08. Fig. 2 shows the distribution of pupal index measurements in households and prediction error mapped across Guayaquil (also see Fig. S3 for corresponding heatmaps and Fig. S4 for relative error). Because of the tightly clustered sampling of some neighborhoods and the contribution of local environmental effects, there was significant spatial autocorrelation (Moran’s I test, p < 0.01). The socioeconomic variables (such as unemployment) captured some of these effects, but there were likely other unmeasured exposures contributing to the spatial pattern. An assessment of multicollinearity, excluding interaction terms, revealed that in our top model, none of the variables tested had variance inflation factor scores above 2, indicating that there is little collinearity between predictors (Table S3).

**Fig. 2.**
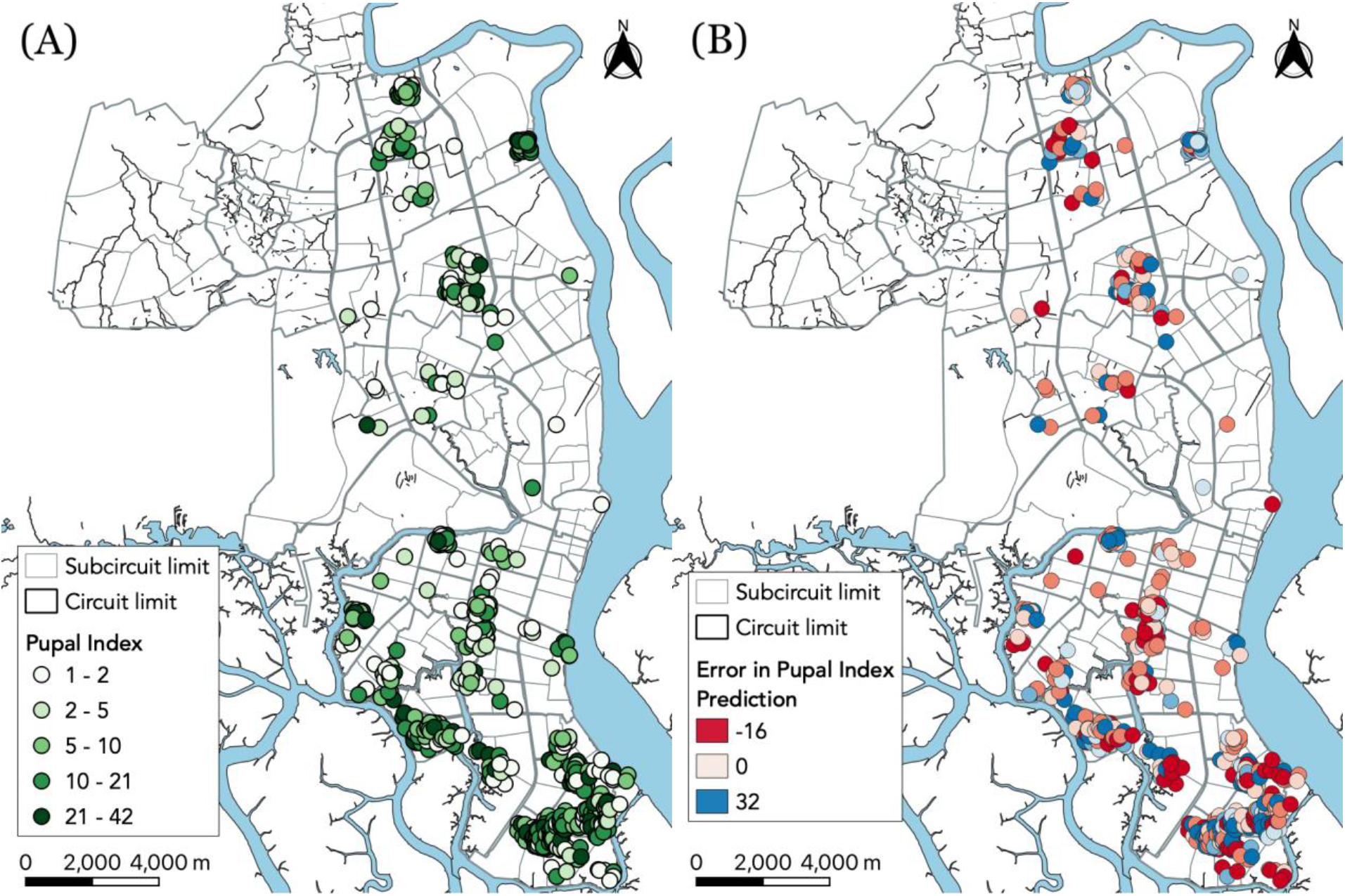
Pupal index measurements (A) and prediction error map (B) based on the final model. Each dot indicates a household included in the final analysis. Error is the difference between the data and the model for each household’s characteristics.

### Influence of container-related factors on Ae. aegypti pupal abundance

We used data from 924 containers to conduct our analysis. The mean pupal sum for the included containers was 15.46, with a standard deviation of 21.63, and a maximum pupal count of 253. We observed that the top container-level models (ΔAICc < 1) consistently selected contaminated water, sewer parts, vases, ceramic material, glass material as appropriate predictors of pupal sum (Table 4, Table S4).

**Table 4.** Variables included in container-level models with the smallest (best) AICc values. Only models that lie within a ΔAICc of 1 of the smallest AICc value are shown. The response variable was the total number of container-level *Ae. aegypti* pupae in each breeding site. Top 5 of all models with ΔAICc < 1 displayed.

| Model | df | LogLik | $\Delta AICc$ |
| --- | --- | --- | --- |
| Car Parts + Contaminated Water + Furniture + Ceramic Material + Glass Material + Metal Material + Plastic Material + Sewer + Vase | 11 | -3459.19 | 0 |
| Car Parts + Contaminated Water + Ceramic Material + Glass Material + Metal Material + Plastic Material + Sewer + Vase | 10 | -3460.23 | 0.03 |
| Bamboo + Car Parts + Contaminated Water + Furniture + Ceramic Material + Glass Material + Metal Material + Plastic Material + Sewer + Vase | 12 | -3458.20 | 0.08 |
| Bamboo + Car Parts + Contaminated Water + Ceramic Material + Glass Material + Metal Material + Plastic Material + Sewer + Vase | 11 | -3459.24 | 0.11 |
| Bucket Part + Car Parts + Contaminated Water + Ceramic Material + Glass Material + Metal Material + Plastic Material + Sewer + Tub + Vase | 12 | -3458.28 | 0.24 |

The average prediction error over 100 simulations of 10-fold cross validation for the container models was 21.54 pupae off of the true mean value of 15.46 pupae. The variables car parts, sewer, vase, ceramic material, glass material, metal material and plastic material were all found to have a significant negative association with the outcome, pupal sum (Table 5). Contaminated water was the singular predictor found to have a significant positive association with pupal sum (Table 5).

**Table 5.** Explanatory variable estimates for the container model with the smallest AICc value. Response variable is pupal sum per container.

| Variable | Log Estimate | Estimate | 95% CI |
| --- | --- | --- | --- |
| Intercept | 3.129 | 22.844 | (17.74, 29.98) |
| Car Parts | -0.666 | 0.513* | (0.35, 0.74) |
| Contaminated Water | 0.247 | 1.280* | (1.10, 1.49) |
| Furniture | -0.484 | 0.616 | (0.35, 1.21) |
| Ceramic Material | -0.942 | 0.390* | (0.23, 0.68) |
| Glass Material | -0.900 | 0.407* | (0.23, 0.74) |
| Metal Material | -0.425 | 0.654* | (0.47, 0.90) |
| Plastic Material | -0.477 | 0.620* | (0.47, 0.81) |
| Sewer | -0.617 | 0.540* | (0.31, 1.02) |
| Vase | -0.414 | 0.661* | (0.46, 0.97) |

\*p < 0.05

## Discussion

The burden of arboviruses transmitted by *Ae. aegypti* in Guayaquil has increased significantly since the 1980s, and targeted interventions are necessary to halt the spread of such diseases [3]. The findings of this study provide evidence that *Ae. aegypti* proliferation is influenced by specific household risk factors. The household models that performed best as determined by AICc contained these variables: canopy use, large solid collection services, unemployment, water volume, water interruptions, biolarvicide, precipitation at week 0, precipitation at week 2 lag, and the interactions: precipitation at week 0 * water interruption and large solid collection services * week 2 lag.

Canopy use has a significant positive association with *Ae. aegypti* abundance, which is counterintuitive and may be attributed to other vector control practices being limited when bed canopies are in use. It is likely that those with higher mosquito populations inside the house are more prone to using bed canopies. The usage of these canopies may result in a false sense of security, or may be an indication of other unmeasured household risk factors that allow for high *Ae. aegypti* densities. The previously mentioned study from Machala, Ecuador also cites an unclear relationship between dengue infections in a city near to Guayaquil, and bed canopy usage [2]. Furthermore, *Ae. aegypti* are daytime feeders so these nets would only affect mosquito feeding if household members are napping during the day.

In our analysis, large solid collection services have a significant negative association with pupal abundance, which may be because there remain fewer untouched breeding sites available for *Ae. aegypti*. Regular bulky item pickup removes tires and other potential mosquito habitats where water could pool. This is corroborated by previous studies which have found that *Ae. aegypti* positive containers were, among the most common, to be trash and flower pots [16].These containers may be more regularly eliminated and maintained in higher socioeconomic areas through large solid collection services. In alignment with this finding, it was also found that unemployment had a significant positive relationship with pupal abundance. This corresponds with the comparison of the socioeconomic status map from Fig. 1, and the pupal index map from Fig. 2, where we find that there is a higher density of *Ae. aegypti* positive households in lower socioeconomic areas. There is also a higher prediction error for these areas as seen in Fig. 2 and Fig. S3, suggesting that areas with higher unemployment need emphasis on research to better understand the specific household risk factors attributed to *Ae. aegypti* pupal density.

Our study also found that average water volume had a significant positive relationship with pupal abundance. A 2012 study from the Tri Nguyen village in Vietnam found that containers where the water volume increased relative to the previous survey had a significantly higher count of *Ae. aegypti* pupae [17]. The study also found that the greatest increase in pupal abundance occurred after a rainfall event. This corresponds to our study’s findings in which both precipitation during week 0 and increasing water volume results in higher APC. Heavy rainfall is known to flush out existing containers, which could explain the negative (not statistically significant) association between rainfall the week of sampling and APC. A negative association (although again not statistically significant) between rainfall with a two-week lag and APC could conceivably result from the same flushing phenomenon, adjusting for the 8-12 days for *Ae. aegypti* eggs to develop into pupae [18,19]. The interaction between large solid collection services and precipitation two weeks prior was significantly positively associated with *Ae. aegypti* pupal counts, which could be due to rainfall providing habitat for eggs to be laid which then develop into pupae 8-12 days later. The meaning of the interaction is somewhat unclear; however, wealthier neighborhoods have increased access to large solid collection services, so there is greater creation and destruction of mosquito habitat compared with poorer neighborhoods. Fig. 2 shows that wealthier areas of Guayaquil have a lower number of high density *Ae. aegypti* households. With fewer breeding habitats in wealthier neighborhoods, precipitation may have a larger and differential effect on pupal density and therefore, any marginal effect may be picked up by the model.

This differentiation is further explained in the water storage practices and distribution of houses. In neighborhoods with higher employment rates and lower illiteracy, there are an increased number of natural areas where precipitation may collect, especially since houses are spread further apart. In less developed neighborhoods, there are different relationships with standing containers. During the rainy season, there are not as many water-holding containers because the water is constantly replenished by the rain. However, in the dry season, there are more standing containers because water is scarcer and needs to be stored for the households.

When the interaction term is included in the final model, precipitation at a week 2 lag has a negative correlation with the outcome. When the interaction term is not being controlled for, 14 days after a precipitation event correlates with higher pupal density. This suggests that there is a specific relationship between large solid collection services and precipitation at a week 2 lag on our outcome, pupal density. However, since large solid services may serve as a proxy for socioeconomic status, this may suggest that there is a dynamic effect across socioeconomic statuses. These nuances are difficult to account for within the model context. For vector control efforts to be effective, it may require a more thorough understanding of the relationship between rainfall and socioeconomic factors that influence pupal density.

*Ae. aegypti* are highly adaptable mosquitoes that were historically found in forested areas using tree holes for breeding but have since adapted to breeding in tires, vases, and other objects found in proximity to human habitations [20]. Their resilience and adaptability pose difficulties when searching for effective control methods, especially for outdoor areas [20]. However, in light of our analyses, certain types and materials of containers may be more or less productive than others.

In our study, vase-type containers were found to be a significant predictor and were correlated with lower pupae counts. Glass material composition was selected as an appropriate explanatory predictor and is correlated with lower pupae counts as well. Vases, other glass-type containers, metal material and ceramic material containers, may have more variable water temperature that impedes *Ae. aegypti* development. Contaminated water was found to be a significant predictor that was correlated with higher pupae counts. The survey had field technicians qualitatively assess if water was contaminated, so it was not quantifiably measured. Research suggests that *Ae. aegypti* prefer “clean water,” but this is a relative designation, as some nutrients in the water may support mosquito populations [7]. Contaminated water may have organic components within the container that promote algal growth and support mosquito proliferation. Water that is contaminated is likely to be untouched and stagnant, allowing *Ae. aegypti* to lay eggs and develop as opposed to cleaner water which may be flushed more often [7]. This finding is corroborated by previous studies that have noted that poor sanitation and water storing habits provide viable habitats for *Ae. aegypti[3]*.

These results suggest that trash collection services targeting large solids, and monitoring of containers that could serve as juvenile mosquito habitat contribute to suppressing *Ae. aegypti* pupal proliferation and consequent adult mosquito densities. These predictive models provide household factors of interest that could be included in future surveys to test hypotheses or assessed in rigorous causal models.

### Limitations and future directions

For the top household model, the mean error was high (10 pupae off of the true value) relative to the mean pupal index (11.0836). However, the standard deviation of the data is 10.263 indicating that the high error is due to the relatively high variance of the data, and the maximum pupal count is 253. Overdispersion and high variance are common in insect count data, therefore these results remain valid [15].

There were months without sampling in each of the years for each of the three parts of Guayaquil; however, they did not share the same months missing in each area, so it was not possible to address this through a time-series analysis to account for the repeated measurements on households. Predictive modeling has limitations. Best subset selection assesses 2^*p*^ models, where *p* indicates the number of parameters, making the implementation of every interaction computationally infeasible when the number of parameters is large. Using previous literature, we assessed the most pertinent interactions and limited our model variable subset selection to 2^21^ models (Fig. S1). This study could be improved with the inclusion of zero APC households to differentiate between containers and households that have zero mosquito pupae compared with those that have positive counts. Additionally, a longitudinal study, as opposed to the cross-sectional study design here, could track temporal dynamics in pupae populations. With a longitudinal study, a time series analysis would be able to assess changing exposures to vector control methods and the environment and any subsequent changes in mosquito populations.

Future studies could correlate pupae counts with household demographics such as age and sex of inhabitants. Noting behavioral differences across these characteristics could also inform efforts for reducing mosquito proliferation and arbovirus spread. Additionally, further studies should compare our estimates of household factors in Guayaquil to those in more rural settings. Household risk factors such as water service interruption and temephos use may have a larger impact in more rural areas, where water interruptions may be more frequent. A similar study placed on an urban-to-rural gradient may help capture these effects. Additionally, dengue serological data could be incorporated to assess correlation between household risk factors and past exposure to dengue, which would be closer to the health endpoint and valuable for the public health sector. Random-effects modeling may further assess our covariates and outcomes with contextual understanding of variable distributions between and within households. Lastly, an understanding of competing dynamics between *Ae. aegypti* and other species of mosquitoes for habitat, breeding, and feeding would provide further context for targeted interventions in areas where multiple species co-exist.

## Conclusion

The results of this study indicate that household factors influenced *Ae. aegypti* pupae proliferation from 2013 to 2016 in Guayaquil, Ecuador. The most notable household-level risk factors for pupae proliferation were the use of bed canopies, unemployment, and water volume in artificial containers, as well as precipitation with two-week lag in conjunction with large solid collection. The positive association of use of bed canopies may indicate areas with high mosquito populations of several species including *Ae. aegypti*. Having access to municipal services like bulky item pickup was protective against mosquito proliferation in households while higher levels of unemployment had the opposite effect indicating that lower socioeconomic neighborhoods have a distinct relationship with *Ae. aegypti* pupal proliferation that requires further exploration. The positive relationship between bed canopy usage and pupal density is counterintuitive, and may be related to a false sense of security or other factors that put households at risk for high mosquito densities. The impact of rainfall on mosquito density is multifaceted as it may both create new mosquito habitat and “wash out” existing habitat. Providing services and social/technical interventions focused on monitoring and eliminating breeding sites may be important for reducing aquatic-stage mosquito densities in households at risk for *Ae. aegypti*-transmitted diseases. The development of *Ae. aegypti* prediction models contributes to public health efforts in Ecuador by providing information to optimize interventions for reducing mosquito densities and preventing dengue outbreaks.

## Declarations

### Ethics approval & Consent to participate

This research is IRB exempt since it involves non-human and animal subjects.

### Consent for publication

Not applicable.

### Availability of data & materials

The datasets generated and analyzed during the current study are not publicly available as they are the property of the Centro de Investigación en Vectores Artrópodos, Instituto Nacional de Investigación en Salud Pública “Dr. Leopoldo Izquieta Pérez”, Quito, Ecuador. Data are however available from the corresponding author on reasonable request.

### Authors’ contributions

TH developed the hypothesis, analyzed and interpreted the entomological data and authored most of the manuscript. TM assisted in the analysis and conception of the research idea, created the geographical figures and was a main contributor in writing the manuscript. KL performed the geographical analyses to utilize for the geospatial figures. PP and VC guided the main data collection for this project and assisted in the conception of research idea as well as provided guidance on interpretation of the results for the paper. JM provided statistical guidance and was also a contributor in writing the manuscript. All authors read and approved the final manuscript.

## Acknowledgements

We thank Dr. Hsiang-Yu Yuan for helpful comments on the statistical aspects of the manuscript.

## Supplementary Material

**Table S1.** Full list of household candidate variables used to find the best model by AICc.

| Full List of Possible Model Variables | Variable Description |
| --- | --- |
| Illiteracy | A relative subcircuit index on illiteracy |
| Unemployment | A relative subcircuit index on unemployment |
| Overcrowding | A relative subcircuit index on overcrowding |
| Number of Children | Number of children (<18 years) residing in the household |
| Number of Adults | Number of adults ( ≥ 18 years) residing in the household |
| Water Interruption | Whether or not the household experience a water service interruption in the last 24 hours |
| Duration of Water Interruption | Duration of the water service interruption |
| Trash Collection Service | Number of times garbage collections occurs at the household per week |
| Large Solid Collection Service | Municipal service of collecting large furniture, tires or other items |
| Sewer Connection | Whether or not there is a sewer connection to the household's waste system |
| Fumigation in Past Weeks | Spraying of deltamethrin, an insecticide applied inside the house in the last 1-4 weeks |
| Abate in Past Weeks | Also known as temephos, an organophosphate larvicide which is applied by 20 grams of granular product per 189 liters |
| Biolarvicide in Past Weeks | *Bacillus thuringiensis* used to target the larval stage of a mosquito |
| Bed Canopy Use | The usage of a canopy over a bed |
| Protective Window/Door Mesh | Mesh present around doors and windows |
| Volume of Breeding Site | Average dimensions of household breeding sites |
| Water Volume of Breeding Site | Total water volume in all household breeding containers divided by the number of breeding sites |
| Current Dengue Case | Household case of dengue at time of interview |
| Dengue in the Last Month | Household case of dengue within the last month at time of interview |
| Precipitation at week 0 | Rainfall during the week of the household interview |
| Week 1 Precipitation Lag | Rainfall during the week prior to the household interview |
| Week 2 Precipitation Lag | Rainfall two weeks prior to the household interview |

**Table S2.**
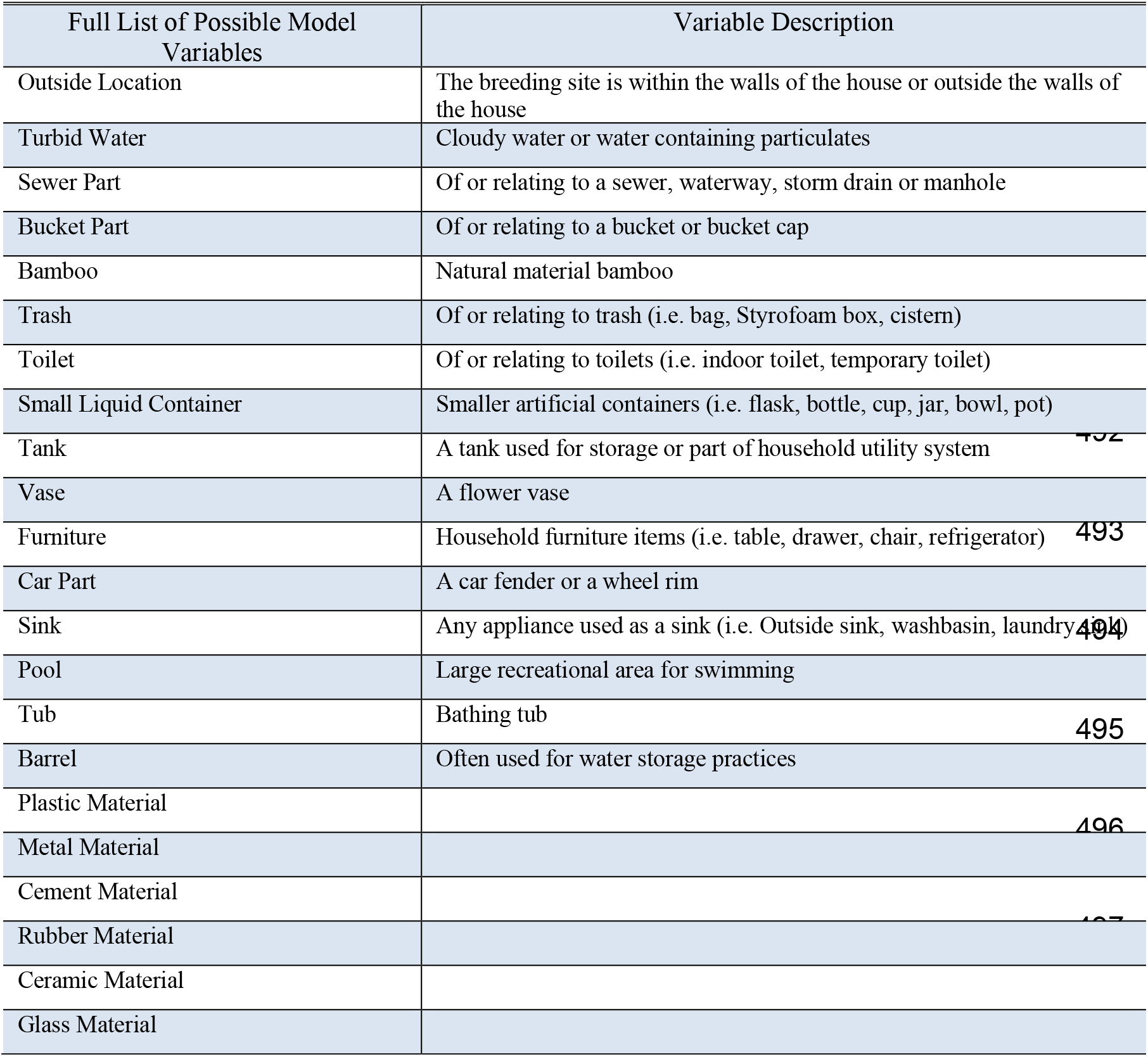
Full list of container-level candidate variables used to find the best model by AICc.

**Table S3.** Variance Inflation Factors (VIF) assess multicollinearity found that (outside of interaction terms) there are no VIF scores >2.

| Variable | VIF |
| --- | --- |
| Large Solid Collection | 1.519 |
| Unemployment | 1.047 |
| Water Volume | 1.250 |
| Canopy Use | 1.519 |
| Precipitation at Week 0 | 1.355 |
| Precipitation at Week 2 Lag | 14.225 |
| Large Solid Collection * Week 2 Lag | 14.612 |

**Table S4.**
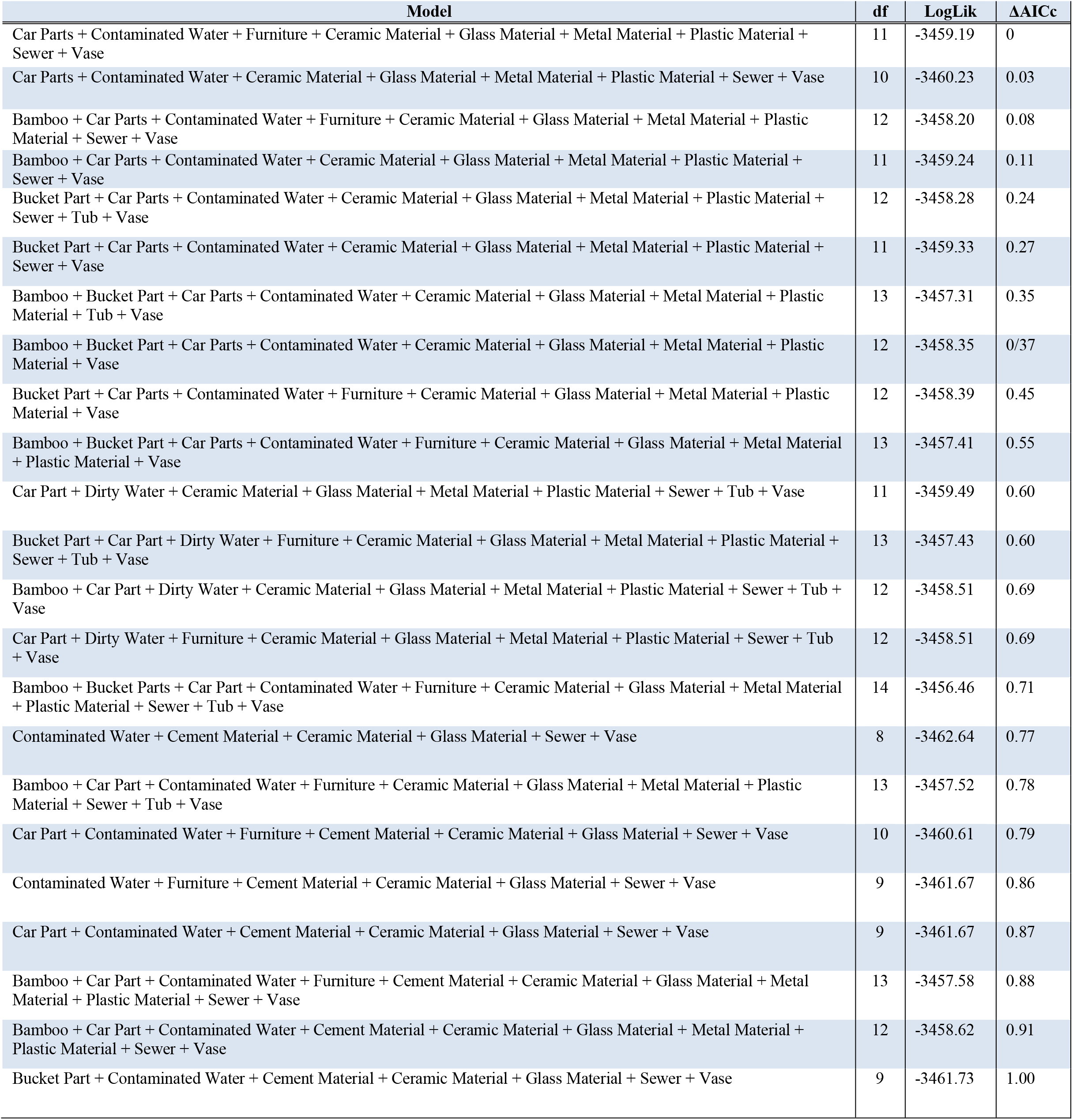
Full list of top candidate models for artificial breeding sites. Top models included have ΔAICc < 1.

**Fig. S1.**
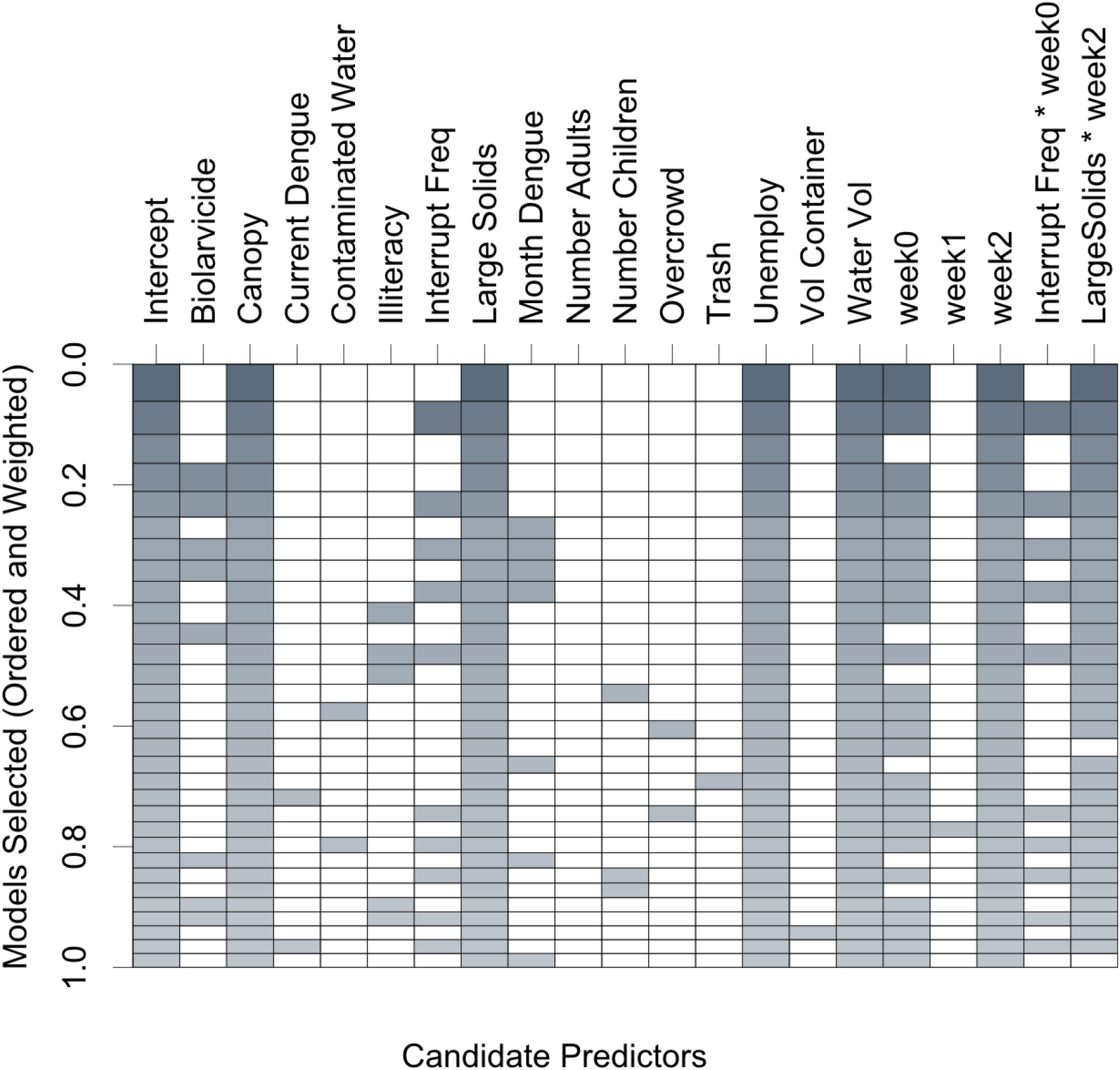
Model selection table where the top axis are all the candidate variables for pupal index (Table S1) and the y-axis represents the frequency the variables were selected in the top candidate models with ΔAICc <2 (31 models shown).

**Fig. S2.**
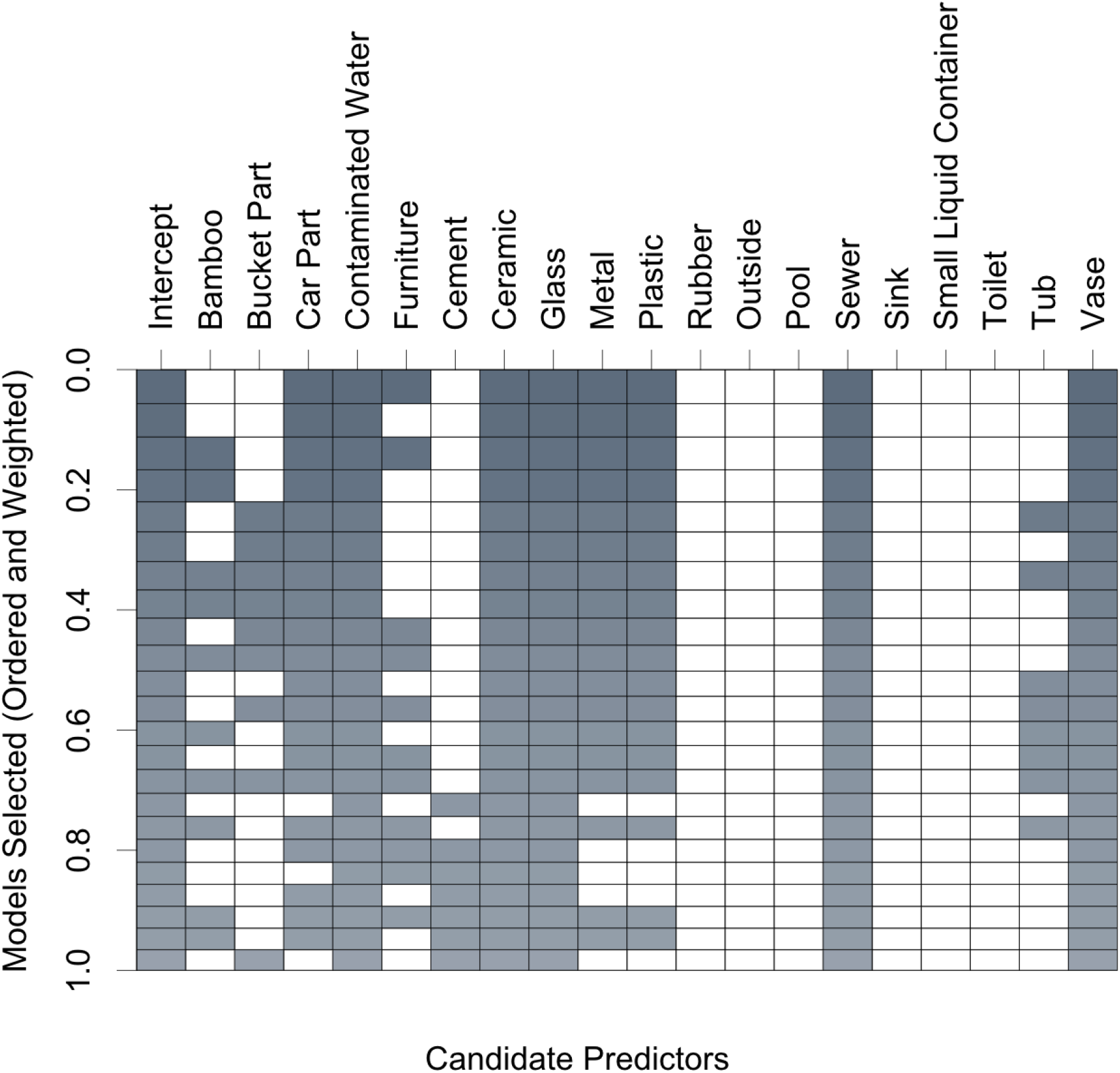
Model selection table where the top axis are all the candidate variables for pupal sum in containers (Table S2) and the y-axis represents the frequency the variables were selected in the top candidate models with ΔAICc < 1 (23 models total shown).

**Fig. S3.**
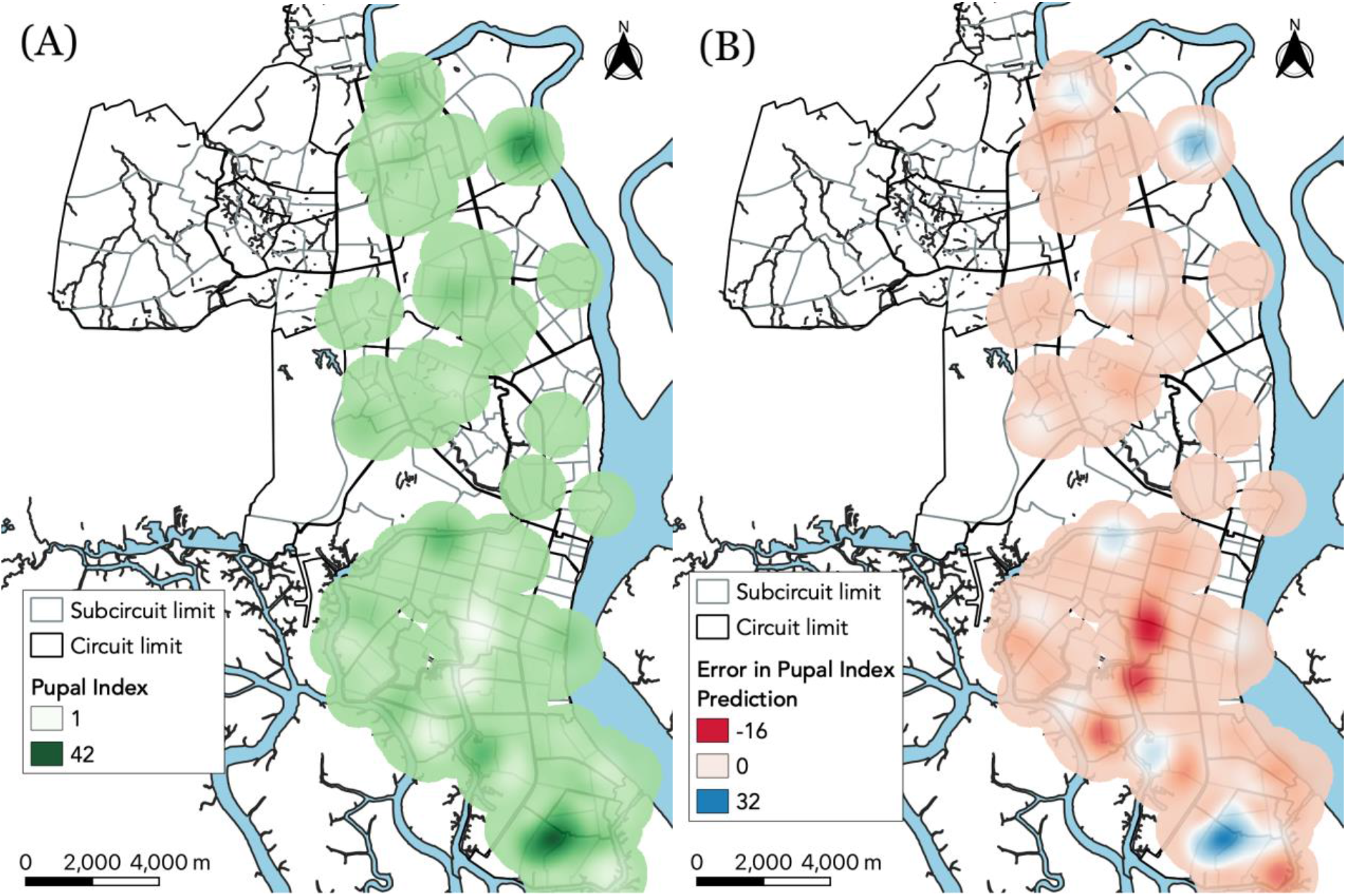
Pupal index measurements (A) and prediction error heatmaps (B) based on the final model. Weighted by value with a bandwidth of 1 km. Error is the difference between the data and the model for each household’s characteristics.

**Fig. S4.**
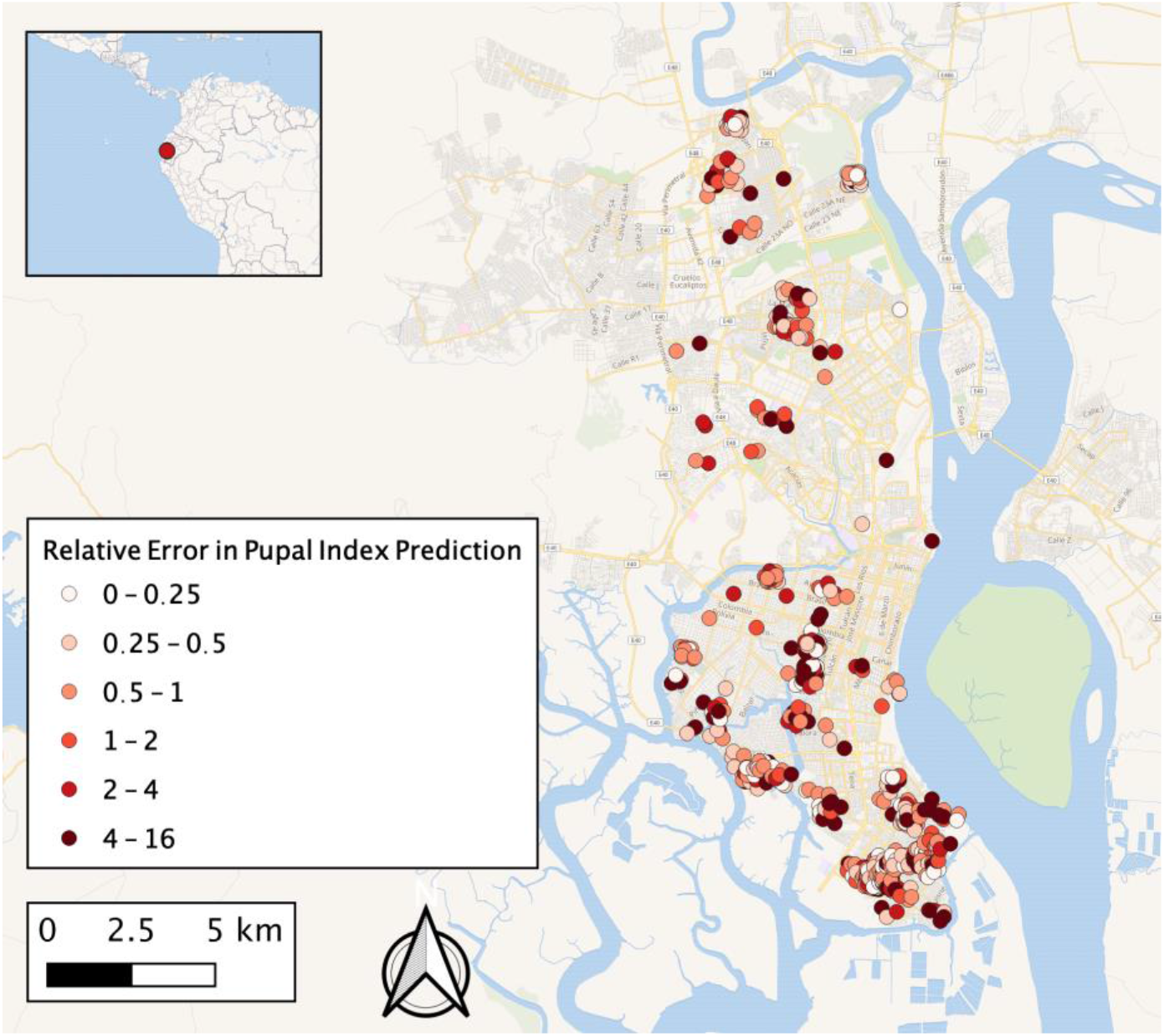
Relative error map for pupal index prediction based on the final model. Each dot indicates the relative prediction error of that household, i.e. the difference between data and model for each household’s characteristics divided by the data.

## References

1. Heydari N, Larsen DA, Neira M, Beltrán Ayala E, Fernandez P, Adrian J, et al. Household Dengue Prevention Interventions, Expenditures, and Barriers to Aedes aegypti Control in Machala, Ecuador. Int J Environ Res Public Health. 2017;14. doi:10.3390/ijerph14020196

2. Kenneson A, Beltrán-Ayala E, Borbor-Cordova MJ, Polhemus ME, Ryan SJ, Endy TP, et al. Social-ecological factors and preventive actions decrease the risk of dengue infection at the household-level: Results from a prospective dengue surveillance study in Machala, Ecuador. PLoS Negl Trop Dis. 2017;11: e0006150.

3. Lippi CA, Stewart-Ibarra AM, Muñoz ÁG, Borbor-Cordova MJ, Mejía R, Rivero K, et al. The Social and Spatial Ecology of Dengue Presence and Burden during an Outbreak in Guayaquil, Ecuador, 2012. Int J Environ Res Public Health. 2018;15. doi:10.3390/ijerph15040827

4. Stewart-Ibarra AM, Muñoz ÁG, Ryan SJ, Ayala EB, Borbor-Cordova MJ, Finkelstein JL, et al. Spatiotemporal clustering, climate periodicity, and social-ecological risk factors for dengue during an outbreak in Machala, Ecuador, in 2010. BMC Infect Dis. 2014;14: 610.

5. Zahouli JBZ, Utzinger J, Adja MA, Müller P, Malone D, Tano Y, et al. Oviposition ecology and species composition of Aedes spp. and Aedes aegypti dynamics in variously urbanized settings in arbovirus foci in southeastern Côte d’Ivoire. Parasites & Vectors. 2016. doi:10.1186/s13071-016-1778-9

6. Ryan SJ, Mundis SJ, Aguirre A, Lippi CA, Beltrán E, Heras F, et al. Seasonal and geographic variation in insecticide resistance in Aedes aegypti in southern Ecuador. PLoS Negl Trop Dis. 2019;13: e0007448.

7. Paul KK, Dhar-Chowdhury P, Haque CE, Al-Amin HM, Goswami DR, Kafi MAH, et al. Risk factors for the presence of dengue vector mosquitoes, and determinants of their prevalence and larval site selection in Dhaka, Bangladesh. PLoS One. 2018;13: e0199457.

8. Abreu FVS de, Morais MM, Ribeiro SP, Eiras ÁE. Influence of breeding site availability on the oviposition behaviour of Aedes aegypti. Mem Inst Oswaldo Cruz. 2015;110: 669–676.

9. Lin C-H, Schiøler KL, Ekstrøm CT, Konradsen F. Location, seasonal, and functional characteristics of water holding containers with juvenile and pupal Aedes aegypti in Southern Taiwan: A cross-sectional study using hurdle model analyses. PLoS Negl Trop Dis. 2018;12: e0006882.

10. Morales D, Ponce P, Cevallos V, Espinosa P, Vaca D, Quezada W. Resistance Status of Aedes aegypti to Deltamethrin, Malathion, and Temephos in Ecuador. Journal of the American Mosquito Control Association. 2019. pp. 113–122. doi:10.2987/19-6831.1

11. George L, Lenhart A, Toledo J, Lazaro A, Han WW, Velayudhan R, et al. Community-Effectiveness of Temephos for Dengue Vector Control: A Systematic Literature Review. PLoS Negl Trop Dis. 2015;9: e0004006.

12. Zhang Y, Ye Z, Lord D. Estimating Dispersion Parameter of Negative Binomial Distribution for Analysis of Crash Data. Transportation Research Record: Journal of the Transportation Research Board. 2007. pp. 15–21. doi:10.3141/2019-03

13. Hastie T, Tibshirani R, Friedman J. The Elements of Statistical Learning: Data Mining, Inference, and Prediction. Springer Science & Business Media; 2013.

14. Burnham KP, Anderson DR. Model Selection and Inference. 1998. doi:10.1007/978-1-4757-2917-7

15. Sileshi G. Selecting the right statistical model for analysis of insect count data by using information theoretic measures. Bull Entomol Res. 2006;96: 479–488.

16. Walker KR, Williamson D, Carrière Y, Reyes-Castro PA, Haenchen S, Hayden MH, et al. Socioeconomic and Human Behavioral Factors Associated With Aedes aegypti (Diptera: Culicidae) Immature Habitat in Tucson, AZ. J Med Entomol. 2018;55: 955–963.

17. Jeffery JAL, Clements ACA, Nguyen YT, Nguyen LH, Tran SH, Le NT, et al. Water level flux in household containers in Vietnam--a key determinant of Aedes aegypti population dynamics. PLoS One. 2012;7: e39067.

18. Eisen L, Monaghan AJ, Lozano-Fuentes S, Steinhoff DF, Hayden MH, Bieringer PE. The Impact of Temperature on the Bionomics ofAedes(Stegomyia)aegypti, With Special Reference to the Cool Geographic Range Margins. Journal of Medical Entomology. 2014. pp. 496–516. doi:10.1603/me13214

19. Tun-Lin W, Burkot TR, Kay BH. Effects of temperature and larval diet on development rates and survival of the dengue vector Aedes aegypti in north Queensland, Australia. Medical and Veterinary Entomology. 2000. pp. 31–37. doi:10.1046/j.1365-2915.2000.00207.x

20. Weaver SC, Reisen WK. Present and future arboviral threats. Antiviral Res. 2010;85: 328–345.

